# Tracking multiple conformations occurring on angstrom-and-millisecond scales in single amino-acid-transporter molecules

**DOI:** 10.1101/2022.07.29.501985

**Authors:** Yufeng Zhou, John H. Lewis, Zhe Lu

## Abstract

The AdiC transporter facilitates the movement of arginine and its metabolite across the membrane of pathogenic enterobacteria, enabling them to evade a host’s highly acidic gastric defense barrier to reach the intestines. Like other transporters, AdiC undergoes a series of necessary conformational changes. Detection of these changes, which occur on angstrom-and- millisecond scales, remains extremely challenging. Here, using a high-resolution polarization-microscopic method, we have successfully resolved AdiC’s four conformations by monitoring the emission-polarization changes of a fluorophore attached to an α-helix that adopts conformation-specific orientations and, furthermore, quantified their probabilities in a series of arginine concentrations. The *K*_D_ values determined for arginine in four individual conformations are statistically comparable to the previously reported overall *K*_D_ determined using isothermal titration calorimetry. This demonstrated strong resolving power of the present polarization-microscopy method will enable an acquisition of the quantitative information required for understanding the expected complex conformational mechanism underlying the transporter’s function, as well as those of other membrane proteins.

## Introduction

Biological membranes enclose individual cells and thereby separate them from their environments. A pure lipid membrane would prevent almost all biologically important substances from effectively getting into or out of cells. The substantial transmembrane movement of these substances are made possible primarily by a class of membrane proteins called transporters. To transport one or more types of substance, a transporter molecule undergoes many necessary conformational changes. An understanding of the mechanism nderlying these changes of a transporter, or other types of protein, requires the ability to resolve individual conformational states. However, three-dimensional protein-conformational changes are not only rapid but also occur often on an angstrom scale. For these reasons, at the single molecule level, reliably resolving the multistate conformational changes of typical proteins, which occur in four dimensions (4D) on angstrom-and-millisecond scales, and acquiring a sufficient amount of data for understanding complex kinetic mechanisms remain extremely challenging.

Importantly, while a multidomain protein molecule undergoes conformational changes, its domains typically rotate relative to one another, in which some secondary structures, e.g. α-helices, inevitably adopt unique spatial orientations in each conformational state. Thus, this feature offers the opportunity to track these conformational states, without the need of determining their detailed three-dimensional (3D) features, by monitoring such an α-helix’s spatial orientation defined in terms of the inclination and rotation angles (*θ* and *φ*; Fig. 1A) with a method of adequate resolution.

**Figure 1.**
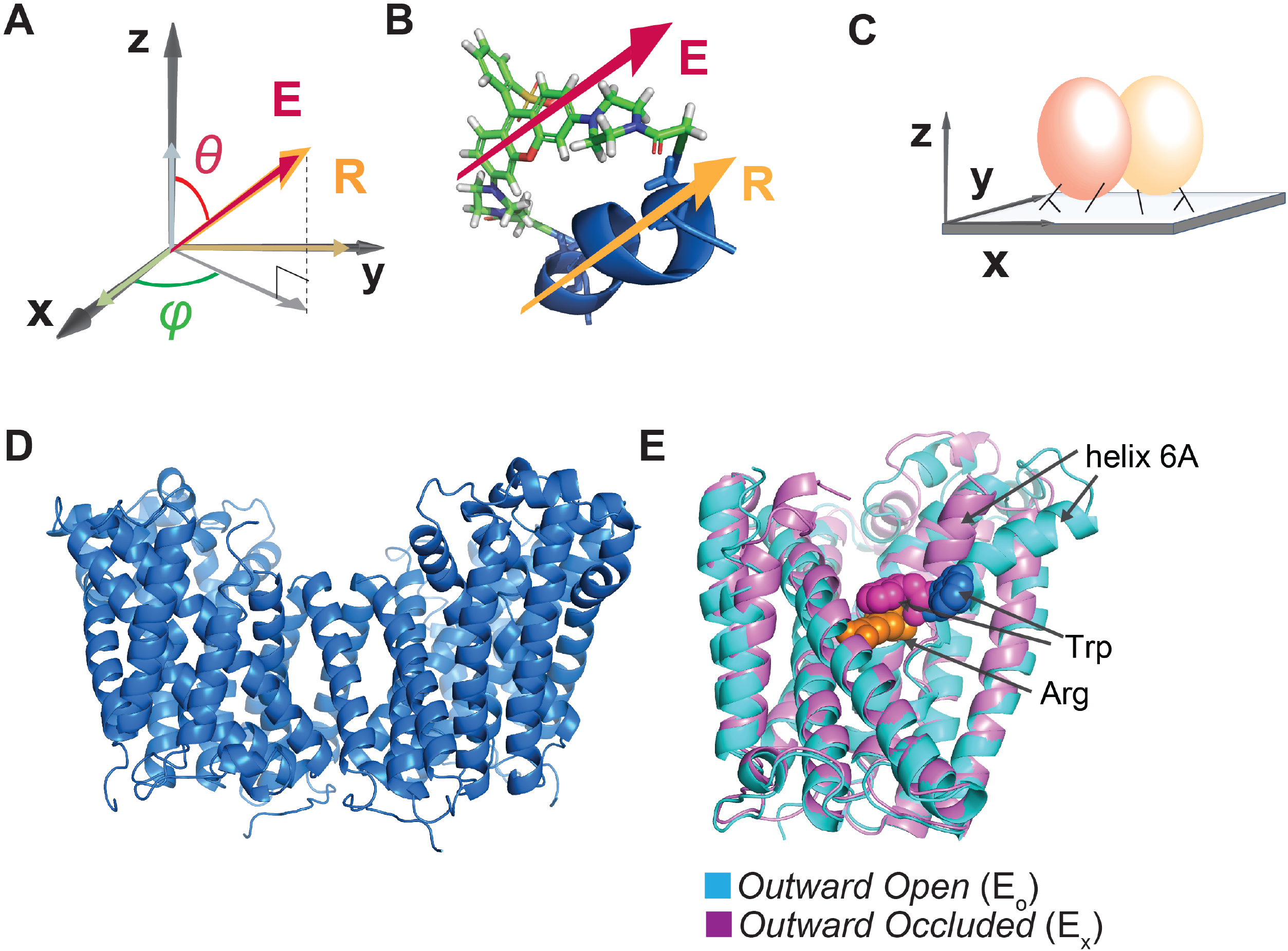
Illustration of the attachment of fluorophore to the AdiC protein and the protein to glass surface. (A and B) The orientation of the fluorophore dipole, defined in in terms of *θ* and *φ*, can be directly related to that of the alpha helix (A), to which bifunctional rhodamine is attached via two mutant cysteine residues (B) where both orientations of the fluorophore dipole and the helix are indicated by the respective arrows. (C) Cartoon illustrating the attachment of an AdiC molecule to a streptavidin-coated coverslip glass via a biotin moiety covalently linked to the N-terminus and two streptavidin-binding tags fused to the N- and C-termini in each of its two subunits, totaling six available sites for binding to streptavidin molecules. D. Structure of AdiC shown as a dimer (PDB: 7O82). (E) Spatially aligned structures of E_O_ and E_X_ states of AdiC shown with a single subunit (PDB: 3OB6, 3L1L). Helix 6, chosen as a labeling site, is indicated in either structure. The substrate Arg^*+*^ (orange) and a Trp residue external to it in the two states (blue and purple) are represented using space-filling models.

One effective way to track the orientation change of an α-helix in a protein domain is to monitor the emission polarization change of a bifunctional rhodamine attached to the helix (Fig. 1B), using a polarization microscope (Sase et al., 1997, Warshaw et al., 1998, Ha et al., 1998, Adachi et al., 2000, Sosa et al., 2001, Forkey et al., 2003, Beausang et al., 2008, Rosenberg et al., 2005, Forkey et al., 2005, Fourkas, 2001, Ohmachi et al., 2012, Lippert et al., 2017, Lewis and Lu, 2019c). The polarization of individual emitted photons, unlike their travel direction, is not meaningfully affected by the diffraction caused by a so-called polarization-preserving objective (Fourkas, 2001). The documented resolutions of such polarization-based detection of rotation motion within a protein had been **≥**25°, estimated on the basis of 2.5 times of the standard deviation (σ) of angle measurements (Rosenberg et al., 2005, Forkey et al., 2005, Ohmachi et al., 2012, Lippert et al., 2017). Recently, our group assembled a polarization microscope with four-polarized-emission recording channels and tracked the orientation change of an isolated, soluble domain of a K^+^ channel via a bi-functional fluorophore attached to an α-helix (Lewis and Lu, 2019c, Lewis and Lu, 2019a, Lewis and Lu, 2019b). By finding optimal hardware, devising necessary numerical corrections for certain system parameters, and developing essential analyses, we have achieved an effective *σ* as low as 2°, translating to 5° resolution for detecting changes in both *θ* and *φ*. For reference, the estimated median radius of proteins is ∼20 Å (Brocchieri and Karlin, 2005, Erickson, 2009), and a rotation of 5° or 10° of a site 20 Å away from the origin would lead to a 1.7 or 3.5 Å change in the chord distance. Thus, the capability to resolve this small angle change allows one to track protein-conformational changes that occur on an angstrom scale in proteins of typical sizes.

Here, we investigate the whole bacterial transporter AdiC protein (Gong et al., 2003, Iyer et al., 2003), which is the first time that this high-resolution fluorescence-polarization-based method is used in the investigation of the conformational mechanism of integral membrane proteins. Our goals are to resolve the multi-state conformational changes, and determine the ligand-dependent energetics underlying these changes. AdiC, a member of the amino-acid and polyamine organocation (APC) transporter superfamily (Jack et al., 2000, Casagrande et al., 2008, Bosshart and Fotiadis, 2019), is a critical component of a system in pathogenic enterobacteria, e.g., *Escherichia, Salmonella*, and *Shigella*, which effectively extrudes intracellular protons, as the bacteria pass a host’s highly acidic gastric defense barrier with a pH value of as low as 2 before reaching the intestines (Gong et al., 2003, Foster, 2004, Fang et al., 2007, Iyer et al., 2003, Krammer and Prevost, 2019). AdiC facilitates the movement of arginine (Arg^+^) into and agmatine (Agm^2+^) out of bacteria, along their gradients. Inside bacteria, Arg^+^ is rapidly decarboxylated to Agm^2+^ by the enzyme AdiA, consuming a proton (Gong et al., 2003, Foster, 2004, Fang et al., 2007, Iyer et al., 2003, Tsai and Miller, 2013). An exchange of extracellular Arg^+^ between a single positive charge and intracellular Agm^2+^ of two charges effectively extrudes H^+^. The direction of the net exchange of the two substrates is dictated by their natural energy gradient. However, as an intrinsic property, AdiC can facilitate the movement of a given substrate in either direction, where the transport of one type of substrate does not markedly depend on which type of substrate is on the opposite side.

## Results

### Fluorescence intensity recordings and transition detection

For microscopic examination, individual purified recombinant AdiC molecules inserted into nanodiscs (Ritchie et al., 2009, Denisov et al., 2019), which were attached to streptavidin adhered to the surface of a piece of coverslip glass via a biotin-moiety covalently linked to the N-terminus and the streptavidin-binding tags linked to the N- and C-termini in each of its two functionally independent and structurally symmetric subunits, totaling 6 sites available for the binding of streptavidin molecules (Fig. 1C; Methods). To examine conformational changes of AdiC, we chose to monitor helix 6a in one of its subunits (Fig. 1D,E) (Gao et al., 2009, Fang et al., 2009, Gao et al., 2010, Kowalczyk et al., 2011, Ilgu et al., 2016, Ilgu et al., 2021). This helix adopts differing spatial orientations among different known structural states in AdiC (Fig. 1E) and other transporters of the same structural fold (Shaffer et al., 2009, Errasti-Murugarren et al., 2019). A bi-functional rhodamine molecule was attached to helix 6a in the region extracellular to the substrate-binding site to avoid affecting the binding affinity (see Discussion). The attachment was through a pair of mutant cysteine residues spaced seven residues apart, aligning the fluorophore dipole along the axis of the helix (Corrie et al., 1998) (Fig. 1B). The density of protein molecules on a cover slip was sufficiently low such that individual fluorescent particles could be readily resolved spatially on microscopic images. The probability of individual AdiC molecules being attached with a single fluorophore was optimized with an empirical protein-to-fluorophore ratio during the labeling procedure, and such molecules were identified during the off-line analysis on the basis of a single-step bleaching of fluorescence. Hereafter, unless specified otherwise, AdiC simply refers to one of its two functionally independent subunits (Fig. 1D,E).

The light emitted from individual attached fluorophores was captured via the objective of a TIRF microscope. To assess the polarization of the captured fluorescence light, we first split it into two equal portions and then further split one portion into 0° and 90° polarized components (*I*_0_ and *I*_90_) and the other portion into 45° and 135° components (*I*_45_ and *I*_135_)(Figs. 2 and 3) (Lewis and Lu, 2019c). These polarized components of fluorescence were recorded with an electron-multiplying charge-coupled device (EMCCD) camera. We integrated the individual intensity images (Fig. 3A) and plotted the resulting values against time. As an example, a set of intensity traces for a molecule examined in the absence or the presence of 0.75 mM Arg^*+*^ is shown in Fig. 3B,C. From these traces, we calculated the trace of total emitted intensity (*I*_tot_) using Eq. 10 (equations with a number greater than 5 are in Methods).

**Figure 2.**
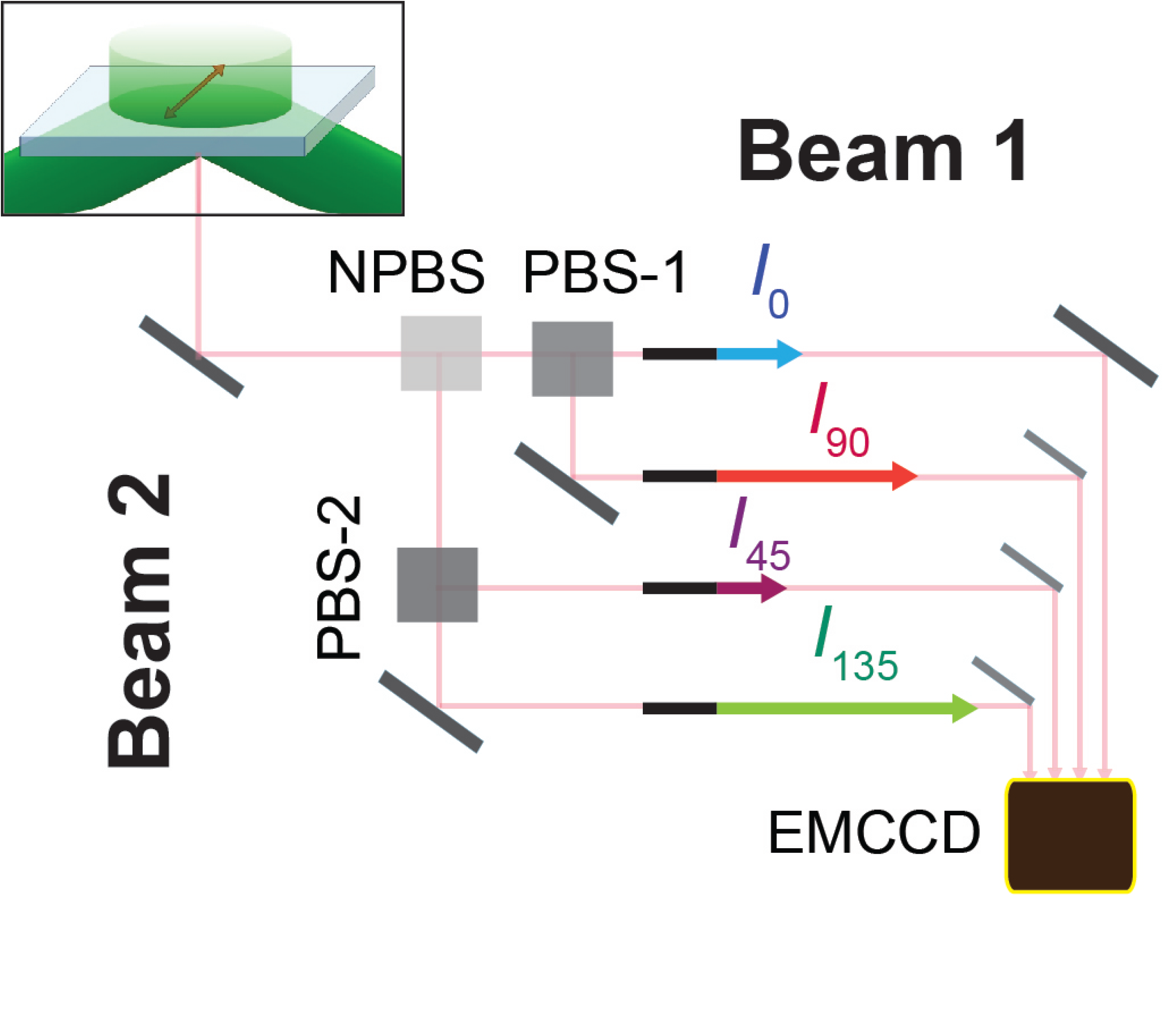
Schematic for four polarized emission intensity collected via a microscope and imaged on an EMCCD camera. Photons, emitted from a fluorophore excited by a circularly polarized laser, are collected by an objective and directed to a non-polarizing beam splitter (NPBS) that splits it eventually to two beams. Beam 1 is further split into its 0 and 90° polarized components (*I*_0_ and *I*_90_) with a glass (N-SF1) polarizing beam splitter (PBS-1), and beam 2 into its 45 and 135° components (I_45_ and I_135_) using a wire grid polarizing beam splitter (PBS-2). These four beams are aligned along one path using pick-off mirrors and directed onto separate sections of an EMCCD camera.

**Figure 3.**
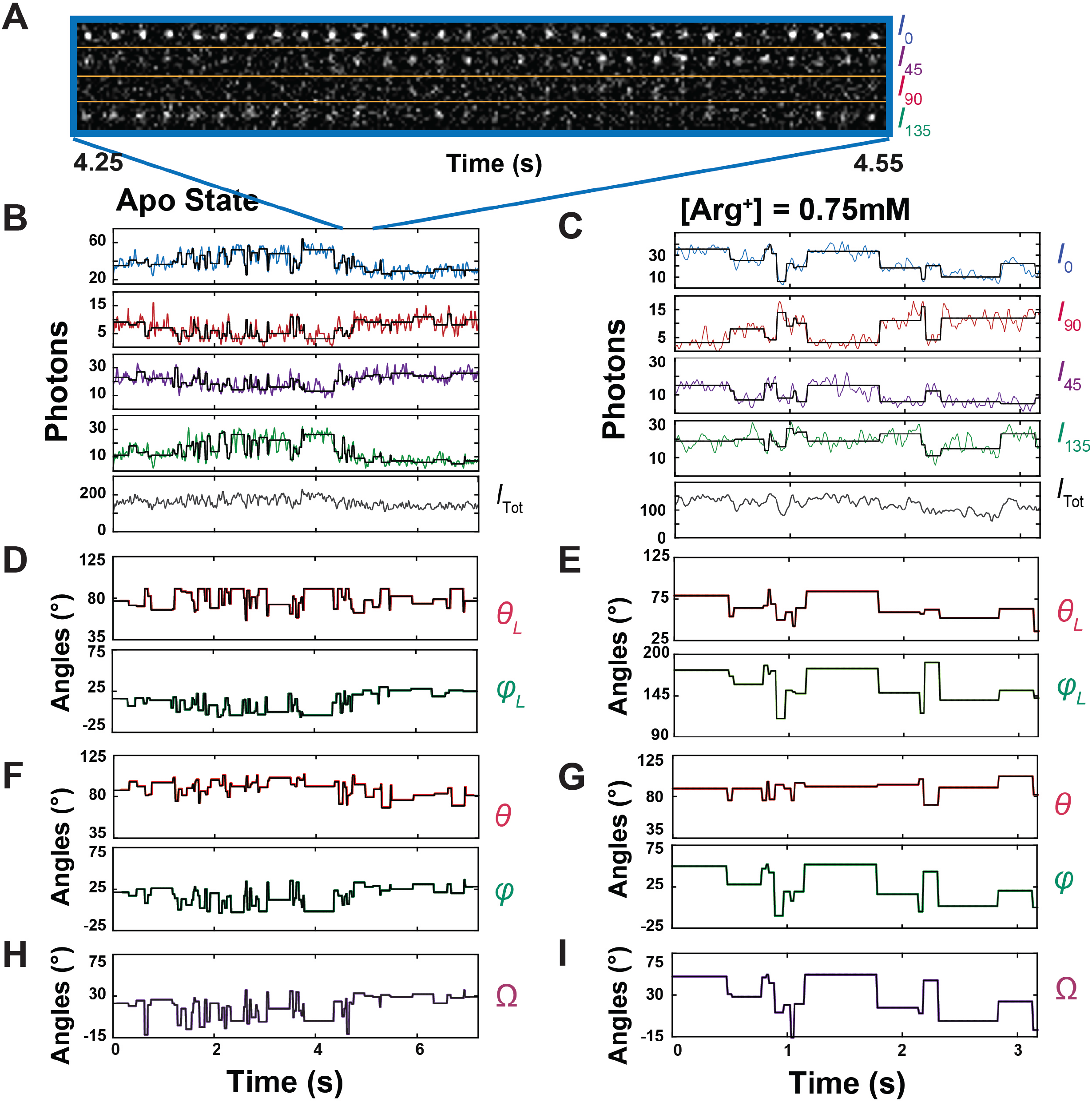
Polarized intensity components of single fluorescent particles and *θ* and *φ* angles calculated from the components. (A) Consecutive frames of four intensity components (*I*_0_, *I*_45_, *I*_90_ and *I*_135_) of a bifunctional-rhodamine-labeled apo AdiC molecule captured over 0 – 300 msec of a 7.2 sec recording (Supplementary Video 1). (B and C) The time courses of integrated intensities color-coded for *I*_0_, *I*_45_, *I*_90_ and *I*_135_ of two bifunctional-rhodamine-labeled AdiC molecules in the absence (B) or presence (**c**) of 0.75 mM Arg^*+*^, from which *I*_tot_ is calculated. Each vertical line in the black traces, superimposed on the colored traces, indicate the time point at which a change in the fluorophore’s orientation is identified, whereas the horizontal lines represent the mean intensity between two identified consecutive time points. (D-G) The traces *θ*_L_ and *φ*_L_ (D and E) in the laboratory frame of reference calculated from black intensity traces, which were rotated into the local frame of refences (F and G). (H and I) Values of *Ω* calculated according to Eq. 31 from the changes in either *θ*_L_ and *φ*_L_ or *θ* and *φ*.

From Eqs. 1 and 2 derived for ideal conditions, one can see that the orientation of the tracked fluorophore would be specifically reflected by the relative intensities, or underlying photon counts, of its four polarized components in the form of a ratio (Lewis and Lu, 2019c) [for the solution of three channels, see reference (Fourkas, 2001)].

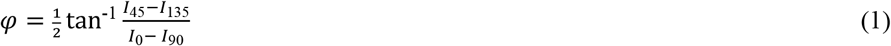

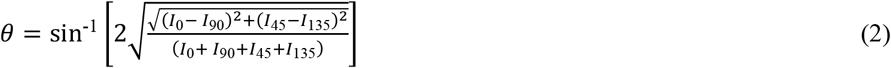

Thus, one can visually notice intramolecular motions that occur on the angstrom-scale from the relative variations in the four intensities (Fig. 3A; Supplementary Video 1).

By examining characteristic changes in the four fluorescence components with a so-called changepoint algorithm (Chen and Gupta, 2001), we detected the time point where a change in the fluorophore’s orientation occurred, which was brought about by the underlying protein conformational change. To do so, we assessed the change in the number of recorded photons (*N*) per unit time with 95% confidence by evaluating the likelihood of two alternative possibilities that a change did or did not occur within a given time interval. This assessment was based on concurrent changes in the recorded *N* among all four polarized components, redundancy that markedly increased the confidence that identified transitions were genuine.

During individual consecutive 10 ms recording intervals, the mean *N* for all four polarization components together was 92 (Methods), with an effective signal to noise ratio (SNR) of 7. As shown in Fig. 3B,C, individual detected transitions are demarcated by the vertical lines in the black traces superimposed on the data traces. From the polarization properties of 92 photons on average, we could reliably resolve the individual transitions in the fluorophore orientation among different conformational states at the intended time resolution. Thus, this method offers an exquisitely sensitive detection of changes in the fluorophore orientation.

### Angle calculation, data averaging and state identification

Resolution of individual conformational states in terms of *θ* and *φ* angles requires a higher SNR and thus a much greater number of photons than what is required for detecting the fluorophore’s orientational changes. One way to solve this problem would be to increase the number of emitted photons by raising the intensity of the excitation laser, but a strong excitation intensity would undesirably shorten the lifetime of the fluorophore. An alternative way to increase SNR is to average the data points within the duration of an event that an examined protein molecule adopts a specific conformation, dubbed dwell time. Our ability to identify individual state-transition points was a prerequisite for us to perform such averaging because individual dwell times were demarcated by these points (Lewis and Lu, 2019c). As shown in Fig. 3B,C, while the vertical lines in the black traces, superimposed on the observed intensities traces, indicate the individual orientation-transition points, the average intensities over individual dwell times are shown as the horizontal lines in the black traces.

From these black traces of *I*_0_, *I*_90_, *I*_45_ and *I*_135_, we calculated *θ*_L_ and *φ*_L_ traces that are color coded (Fig. 3D,E); these two angles are defined in the standard laboratory (L) frame of reference for microscopic studies where the z-axis is defined as being parallel to the optical axis of the objective and the x-y plane parallel to the sample coverslip. All angle calculations were done using expanded versions of Eqs. 1 and 2 (Eqs. 9 and 11), which contain four necessary, pre-determined system parameters. These previously established parameters numerically correct for incomplete photon collection (*α*) (Axelrod, 1979), depolarization caused by imperfect extinction ratios of the polarizers (*f*) (Lewis and Lu, 2019c) and fast wobble motion of the fluorophore dipole (*δ*) (Forkey et al., 2005), and slightly different intensity-recording efficiency of the four polarization channels (*g*). In the angle calculation, we used the information from all photons (980 photons on average) recorded from all four polarization channels during individual dwell times to determine the corresponding (mean) angles over these individual durations with adequately low *σ*. The relation between *σ* for the two angles and SNR among individual analyzed particles is shown in Fig. 4A,B, in which the highest value of <4° translated to a resolution of <10°.

**Figure 4.**
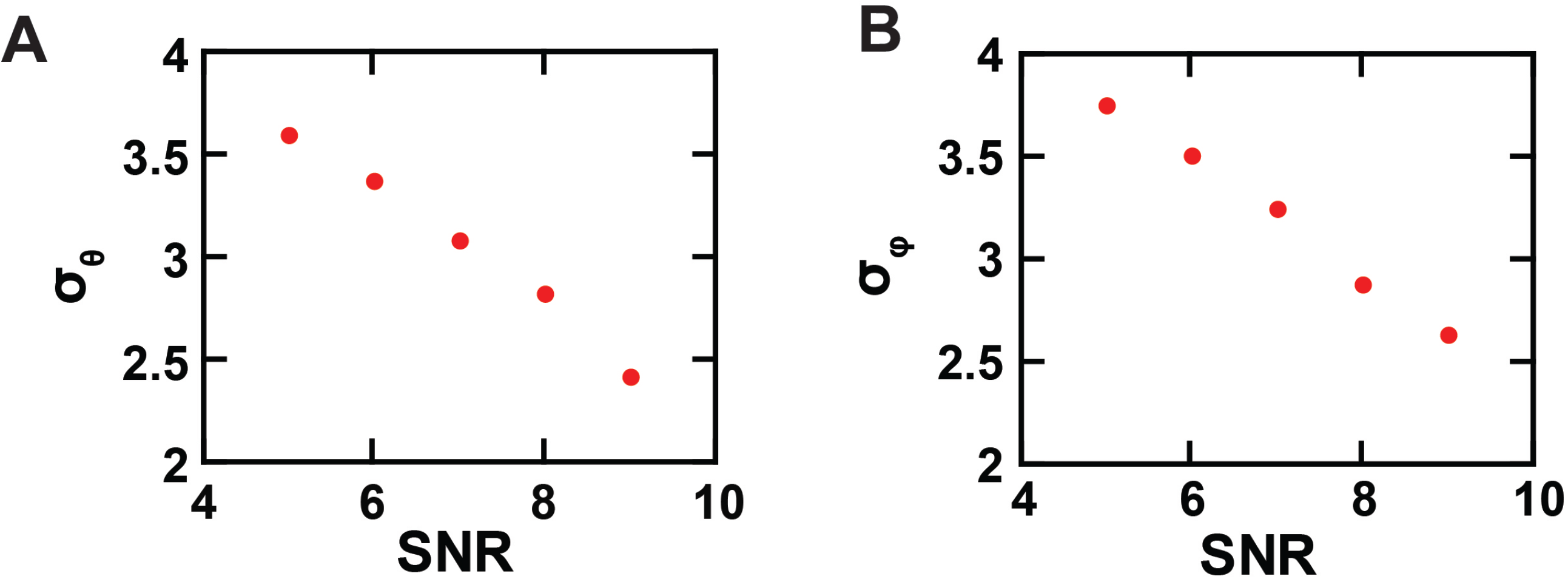
Experimental resolution of conformational states and state-dependent relations of *θ* and *φ*. (A and B) The *σ* value of the *θ* (A) and *φ* (B) populations, built with data acquired from individual single AdiC molecules labelled with bifunctional rhodamine, are plotted against SNR.

Subsequently, conformational state populations were identified from both angles together to increase resolvability and confidence. We sorted the states adopted during individual dwell times into distinct populations using the *k*-means clustering algorithm optimized with two coupled algorithms (*simulated annealing* plus *Nelder-Mead downhill simplex*)(Press et al., 2007). Through this sorting process, we could resolve, from the angle traces as shown in Fig. 3D,E, four conformational state populations, *C*_1_, *C*_2_, *C*_3_ and *C*_4_, ranked according to the values of both angles (Fig. 5A,B or 5C,D; see below for their distinction). When two consecutive events were determined to belong to the same state distribution, the transition between them identified by the changepoint algorithm would be part of the expected false positive outcomes. These events would then be merged to form a single event for the state distribution. This operation reduces the false positive identification of transition points.

**Figure 5.**
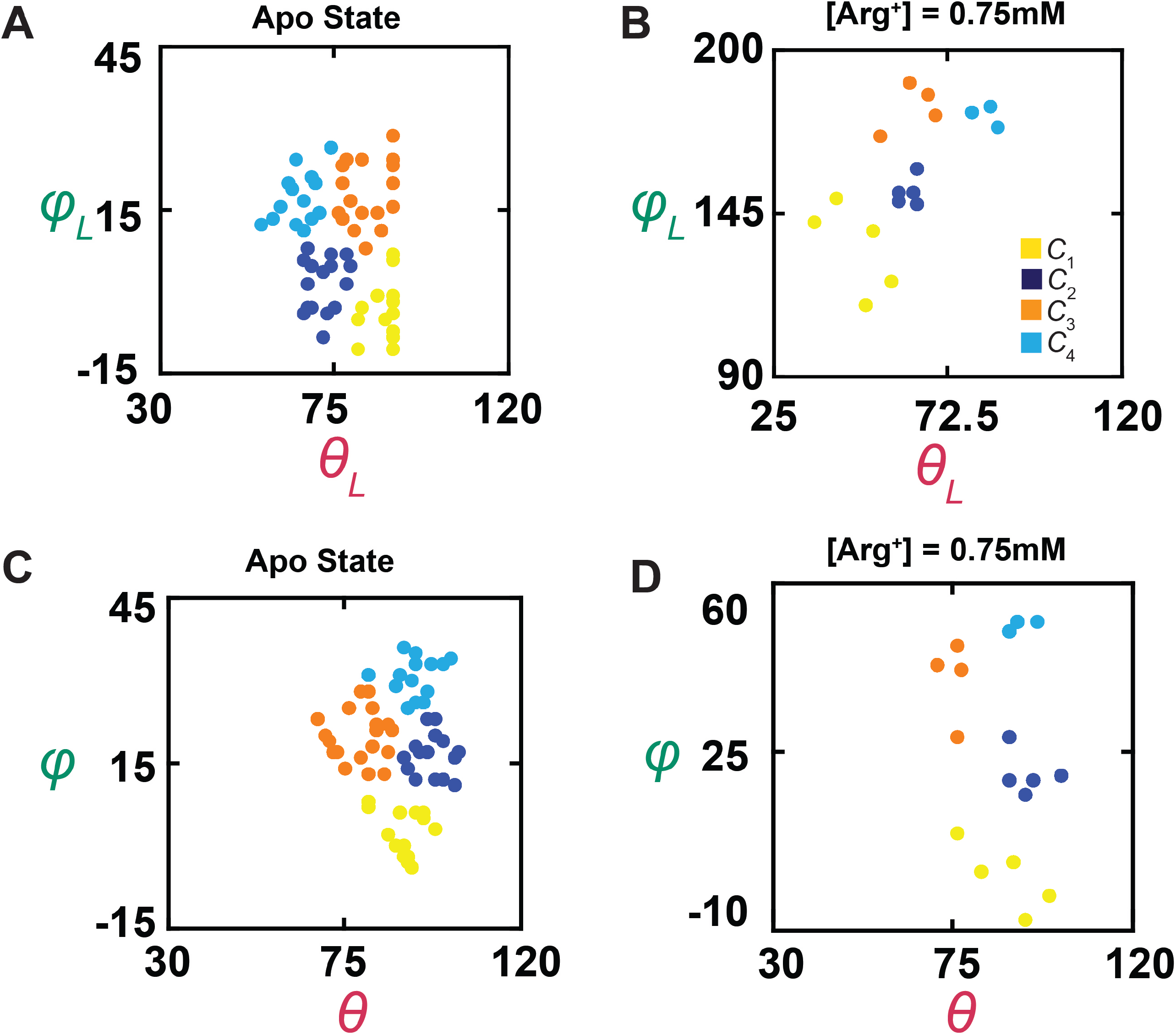
Plots of *θ* versus *φ*. (A-D) *θ* and *φ*, determined in Fig. 3, are plotted against each other in the laboratory (A, B) or the local (C, D) frame of reference as defined in the text, in which the data points for conformational state *C*_1_ are colored yellow, *C*_2_ colored blue, *C*_3_ colored orange, and *C*_4_ color cyan.

Fig. 6 illustrates four superimposed partial structures of AdiC and related transporters (Shaffer et al., 2009, Gao et al., 2010, Errasti-Murugarren et al., 2019, Ilgu et al., 2021), in which the orientation of helix 6 or its counterpart shown in Fig. 6A,B is represented by a vector that is color-coded for a specific structural state (Fig. 6C), with six direct angle *Ω*_i,j_ among four states; all of the vectors are calculated from the corresponding and well-resolved regions. Following identification of four conformational states in the present polarization study, *Ω*_i,j_ between a given reference state *i* and another conformational state *j* was calculated using Eq. 31 from *θ* and *φ* of states *i* and *j* (Fig. 3F,G where *C*_1_ is used as the reference).

**Figure 6.**
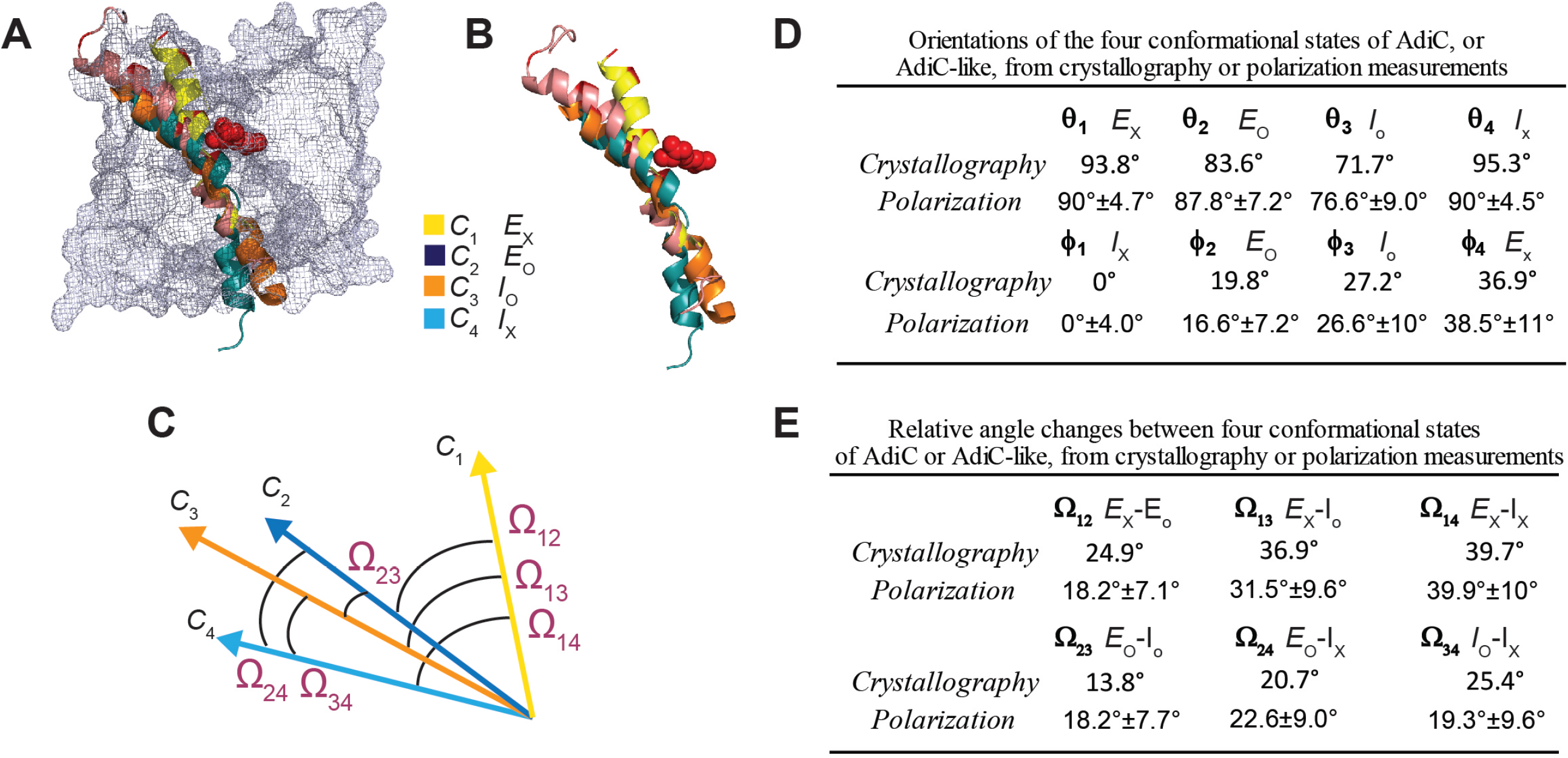
Comparison of the orientations of helix 6A in the corresponding states determined from the crystal structures and in the polarization study. (A and B) Alignments of AdiC helix 6A in the structural states E_O_ (blue)(PDB: 7O82) and E_X_ (yellow)(PDB: 3L1L) with the corresponding helices of BasC and ApcT in the states I_O_ (orange)(PDB: 6F2G) and I_X_ (cyan)(PDB: 3GIA), with (A) or without (B) the rest of the protein represented by a mesh contour. (C) Depiction of the six *Ω* angles among the four orientations of the compared helices in the four states, represented by four arrows color-coded for states. (D, E) The *θ* and *φ* angles (D) of the helix for corresponding states determined from the crystal structures and in the polarization study, as well as the *Ω* angles (E), are compared in the local frame of reference. All angle values for the conformational states determined by polarization are presented as mean ± standard deviation (σ).

### Spatial alignment of individual molecules

For greater likelihood to accurately estimate angle values, we need to determine their mean values from numerous molecules that had already been analyzed individually for necessary spatial resolution. This operation requires all molecules be aligned in the same orientation.

Unfortunately, individual molecules on the coverslip, and consequently the tracked helix, were randomly oriented in x-y plane relative to the *x*-axis, i.e., their *φ*_L_ varying randomly among different molecules. This problem previously prevented us from building the distribution of *φ*_L_ for a given state among individual molecules (Lewis and Lu, 2019c). Furthermore, each dimer molecule of AdiC is anchored to the cover slip coated with streptavidin through two available biotin moieties and four streptavidin-binding tags, each of which was covalently linked to an N-or C-terminus of the polypeptides of two subunits. These terminal binding regions were of some flexibility. Furthermore, the individual dimer molecules have only two-fold symmetry. Given these factors, the two-fold symmetry axis of the individual dimer molecules may not be adequately aligned with the optical (z) axis of the microscope framework. Consequently, the orientation of one molecule could vary considerably from that of another (Fig. 5A,B). Resolving this issue would also pave the way for studying individual randomly oriented molecules in the future.

To align all the molecules during analysis, we mathematically rotate them from the laboratory frame of reference into a local coordinate system defined on the basis of the spatial features of the tracked helix in the protein (supplementary Fig. S1). This system is defined such that the tracked helix in *C*_1_ is always aligned with the local x-axis, i.e., mean *φ*_1_ = 0°, and the helix in the plane defined by *C*_1_ and *C*_4_ is always in the local x-y plane, i.e., mean *θ*_1_ or *θ*_4_ = 90°. These two features fully define the x,y,z-axes, and thus *θ* and *φ* (without subscript notation) in the local framework of AdiC. Evaluated in this local framework (Figs. 3F,G, 5C,D and 8), individual events of the same state observed with all molecules under a given condition could be used to build a single distribution (Fig. 7A,B). From the distribution for each state, we determined the mean *θ*_i_ and *φ*_i_ which are summarized in Fig. 6D (filled circles). Furthermore, we used four unit-vectors with filled heads to specify the tracked helix’s orientations in the four states in the local framework (Fig. 8A), defined by the *θ*_i_ and *φ*_i_ values summarized in Fig, 6D and plotted in Fig. 8B. From the four sets of *θ*_i_ and *φ*_i_ we calculated the six mean direct-angle *Ω*_i,j_ values among the four vectors (Fig. 6E).

**Figure 7.**
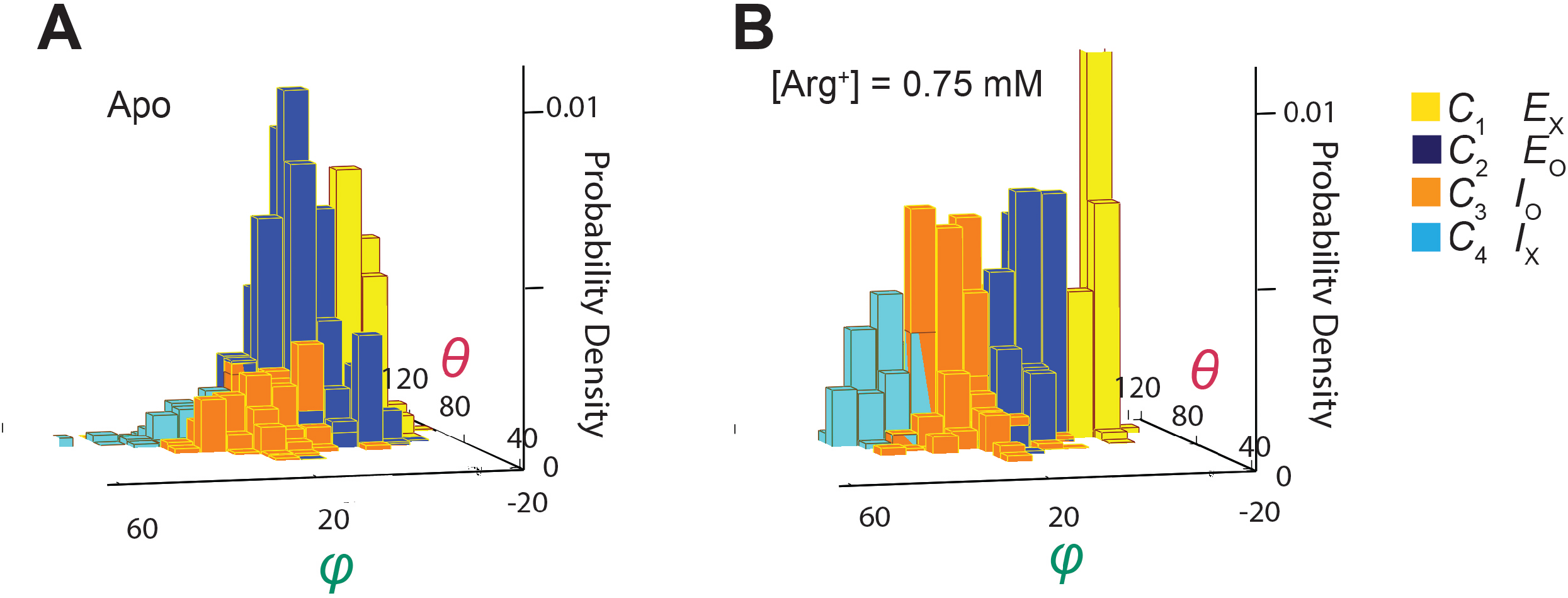
Ensemble 3D probability density distributions of *θ* and *φ*. (A and B) The *θ* and *φ* distributions in the absence (A) or presence of 0.75 mM Arg^*+*^ (B), where the value of *θ* is plotted along the x-axis, the value of *φ* along the y-axis, and the value of probability density along the z-axis. Distributions were built with the data analyzed from 91 or 75 number of particles with a total 3048 or 1494 number of events. Data columns for the conformational state *C*_1_ is colored yellow, *C*_2_ colored blue, *C*_3_ colored orange, and *C*_4_ color cyan.

**Figure 8.**
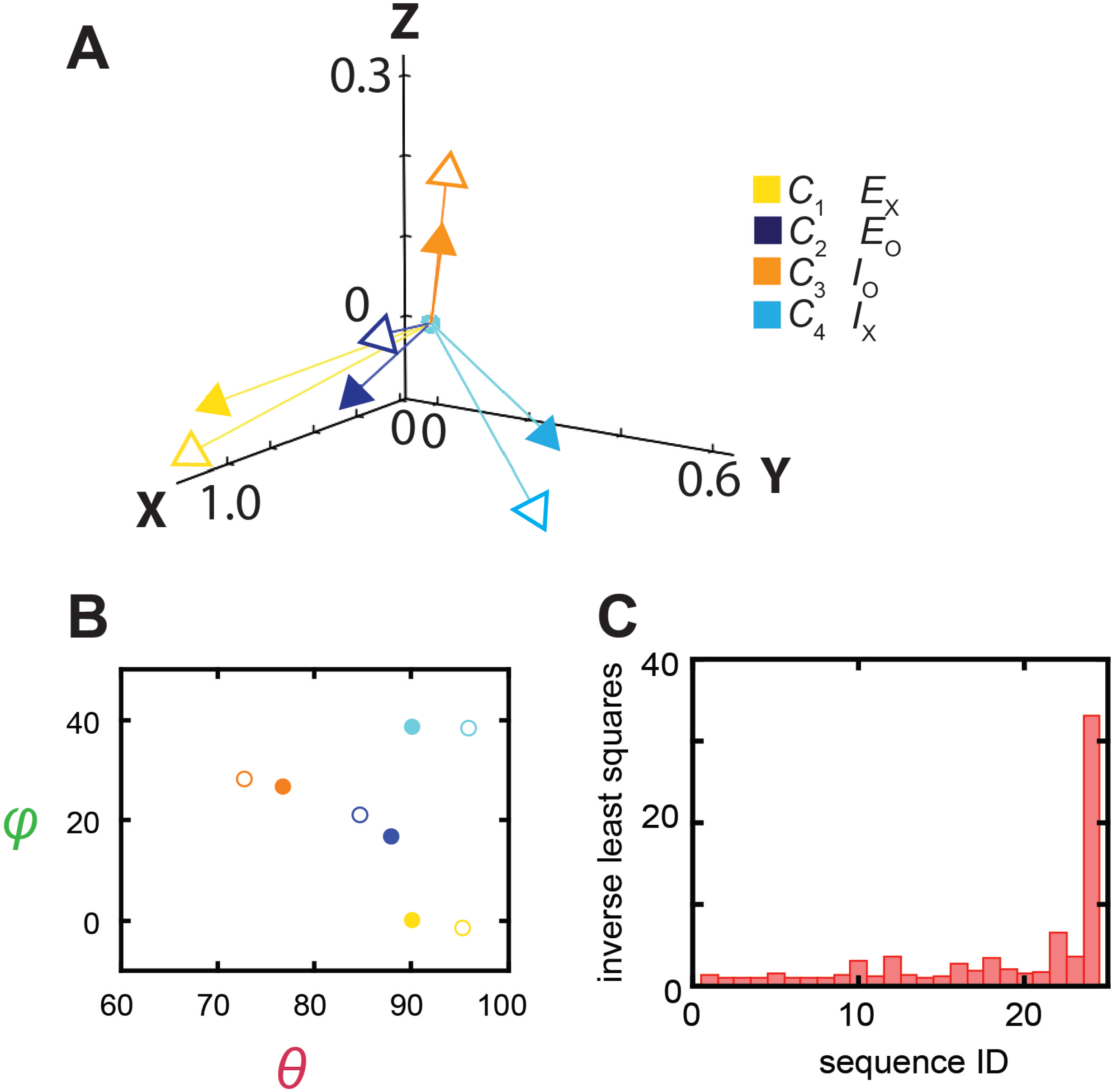
Relations between the structural and conformational states determined respectively from the crystal structures and in the polarization study. (A) The four mean orientations of the helix in the four conformational states are represented by a set of four unit-vectors (closed heads) in the local framework of coordinates whereas those for the four structural states by another set of unit vectors (open heads). The vectors are drawn according to *θ* and *φ* values obtained from the distributions, and color-coded for the corresponding states. The two sets of vectors are overlaid as described in the text. (B) Scatter plots of mean *θ* versus *φ* values (closed circles) for four conformational states, which are compared with those for the four structural states (open circles); all are color coded for states. (C) The inverse values of combined least-distance-squares between the locations of the arrow heads of the two compared groups (open versus closed) in B for all 24 possible combinations among them.

Although the ensemble mean angle values should be more accurate statistically, the apparent angle resolution would be poorer. Indeed, compared with *σ* for individual particles (Fig. 4), the ensemble angle distributions built with data determined from molecule-by-molecule analyses have much larger *σ* (8-11**°**, Figs. 6D, 7A and 7B), except for those of *φ*_1_ and *θ*_1_ or *θ*_4_ which are normalized such that their mean values equal 0° and 90°. Thus, the initial molecule-by-molecule data analysis, as shown in Fig. 3, was essential for resolving individual conformational states in terms of *θ* and *φ*.

### Determination of equilibrium constants among the Apo and Arg^+^-bound conformational states

As an essential evaluation of the basic energetic properties of the protein’s individual states, we examined individual AdiC molecules in the presence of a large series of Arg^*+*^ concentrations, an examination that also helps to relate the states identified here to those structurally and functionally defined states (see Discussion). Both sides of an AdiC molecule facing the same solution allowed us to determine the probabilities of individual conformational states under conditions where the system as a whole is in equilibrium, through which we could straightforwardly obtain equilibrium constants (*K*_i,j_) among the states, including dissociation constants (*K*_*D*i_) to be compared in Discussion with previously reported *K*_*D*_ values.

Shown in Fig. 7A,B is a pair of three-dimensional plots of the probability density distributions of *θ*_i_ and *φ*_i_ under 0 or 0.75 mM Arg^*+*^ conditions, built with the data from numerous individually analyzed particles. From the angle distributions like these two, we obtained the probabilities of occupying each of the four states in a series of Arg^*+*^ concentration (Fig. 9A). All four states appeared in the absence and the presence of Arg^*+*^. With increasing concentration, the probability of *C*_3_ increased whereas that of *C*_2_ decreased. In contrast, the probability of *C*_1_ or *C*_4_ changed only slightly. Nonetheless, these observations indicate that all states bind Arg^*+*^ because any state that could not bind Arg^*+*^ would practically vanish in a saturating concentration of Arg^*+*^. Here, a model with a minimum number of eight states is required to account for the conformational behavior of AdiC: a set of four without ligand bound (*C*_1_, *C*_2_, *C*_3_ and *C*_4_) and another set of four with the Arg^+^ ligand (L) bound (*C*_1_.L, *C*_2_.L, *C*_3_.L and *C*_4_.L) (Fig. 9B). The probability *p*_i_ of occupying state *i* is expressed as:

**Figure 9.**
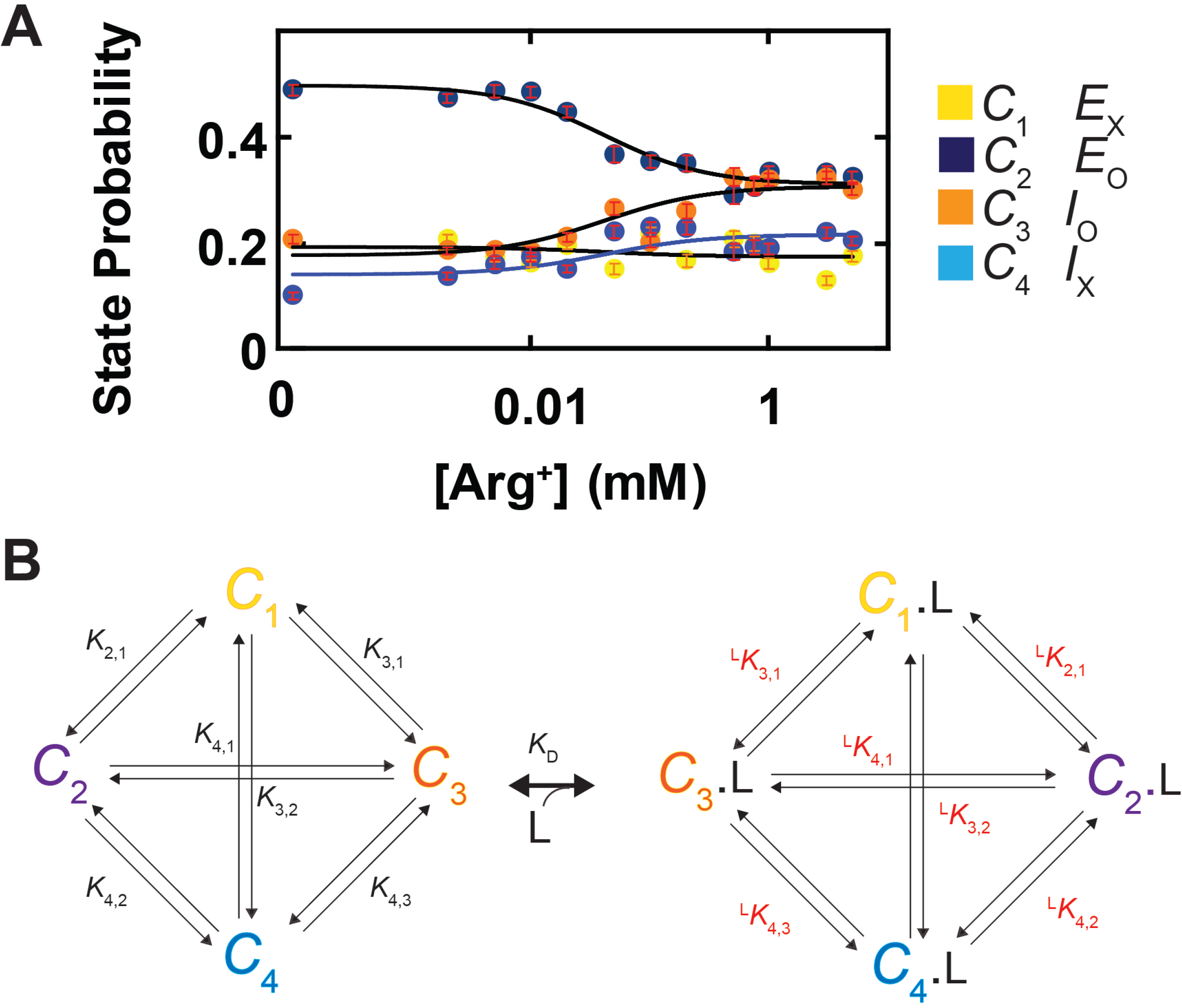
Ligand dependence of the probabilities of conformational states and the diagram of a conformational state model of AdiC. (A) The probabilities of individual states (*C*_1_-*C*_4_) are plotted against the Arg^*+*^ concentration on a logarithm scale. The curve superimposed on the data corresponds to a global fit of a model in which the interaction between the subunit of AdiC and Arg^+^ has a one-to-one stoichiometry. The fitted values of all parameters are summarized in supplementary Table 1. (B) An eight-state model that accounts for the conformational behaviors of AdiC: four apo states and four ligand-bound states.

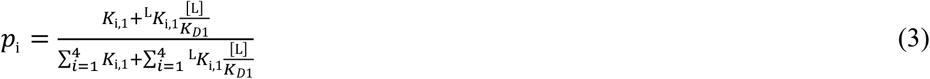

where *C*_1_ is used as a reference for the other states (*C*_i_) to define equilibrium constants:

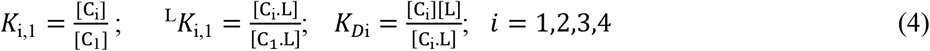

where *K*_1,1_ is defined to equal one. In principle, *K*_i,1_ or ^L^*K*_i,1_ are constrained by the state probabilities under the condition of zero or saturating Arg^*+*^, and *K*_Di_ by the so-called midpoint positions of the curves (Fig. 9A). In practice, we determined these parameters by simultaneously fitting Eq. 3 to the four plots in Fig. 9A, all summarized in Table 1. Together, these constants would fully define the eight-state model quantitatively.

**Table 1:** Equilibrium and dissociation constants for apo and Arg^+^ bound states

| apo state<sup>a</sup> |  |  |  |  |  |
| --- | --- | --- | --- | --- | --- |
| | | $P_1$ | 0.192 +0.015/-0.024 | | |
| $K_{2,1}$ | 2.566 +0.589/-0.244 | $P_2$ | 0.492 +0.040/-0.014 | | |
| $K_{3,1}$ | 0.923 +0.061/-0.149 | $P_3$ | 0.177 +0.011/-0.037 | | |
| $K_{4,1}$ | 0.729 +0.225/-0.103 | $P_4$ | 0.140 +0.026/-0.014 | | |
| Arg<sup>+</sup> bound state |  |  |  | K<sub>D</sub> (μM) |  |
| | | $^{\text{arg}}P_1$ | 0.172 +0.020/-0.020 | $^{\text{arg}}K_{D1}$ | 49 +46/-25 |
| $^{\text{arg}}K_{2,1}$ | 1.795 +0.298/-0.319 | $^{\text{arg}}P_2$ | 0.308 +0.013/-0.025 | $^{\text{arg}}K_{D2}$ | 69 +67/-33 |
| $^{\text{arg}}K_{3,1}$ | 1.782 +0.272/-0.193 | $^{\text{arg}}P_3$ | 0.306 +0.019/-0.018 | $^{\text{arg}}K_{D3}$ | 25 +20/-14 |
| $^{\text{arg}}K_{4,1}$ | 1.250 +0.228/-0.188 | $^{\text{arg}}P_4$ | 0.214 +0.018/-0.015 | $^{\text{arg}}K_{D4}$ | 28 +31/-14 |

## Discussion

Like other transporters, AdiC is expected to adopt four main types of structure-function states in terms of the accessibility of its external and internal sides to substrates, dubbed the externally open (E_o_), externally occluded (E_x_), internally open (I_o_), and internally occluded (I_x_) states (Post et al., 1972, Gao et al., 2010, Krammer and Prevost, 2019). By this definition, when a transport molecule adopts the E_o_ and I_o_ states, it is accessible only to extracellular and intracellular ligands, respectively. The molecule in the E_x_ or I_x_ state is inaccessible to ligands from either side. Thus far, the crystal structures of AdiC in the E_o_ and E_x_ state have been solved (Gao et al., 2009, Fang et al., 2009, Gao et al., 2010, Kowalczyk et al., 2011, Ilgu et al., 2016, Ilgu et al., 2021). Additionally, the structures of BasC and ApcT (Shaffer et al., 2009, Errasti-Murugarren et al., 2019), which have the same folds as AdiC, were solved in I_o_ and I_x_ states, respectively.

For reference, we evaluated the orientation of helix 6 in two AdiC structures and that of its counterparts in the two other structurally related transporters, each of which is in one of the four different types of state (Fig. 6A,B). On the basis of the well-resolved regions of the compared helices in the four structures, the difference in *θ* and *φ*, at least one of them, between each compared pair of helices is >10º. On the basis of this structurally predicted requirement of 10º resolution, we chose to use the intensity data with SNR of ≥ 5, which corresponds to σ for the *θ* and *φ* distributions of ≤ 4º, translated to a resolution of ≤ 10º (Fig. 4). Supporting this choice of the cutoff criterion for SNR, the mean value determined from polarization measurement for the smallest angle change required to be resolved, namely, the difference between *θ*_2_ and *θ*_3_, is ∼12º (Fig. 6D). Such a resolution is within the limit of ≤ 10º set above as σ ≤ 4º.

Next, we examine how these four structural states are related to those states identified in the present polarization study, the latter of which will be referred to as the conformational states for clarity. The total combination between the four structural states and four conformational states yields 24 possibilities. To determine the most probable combination, we fit the unit-vectors (open heads, Fig. 8A) and consequently their *θ* and *φ* (open circles, Fig. 8B), which specify the helix’s orientations in the four structural states, to those for conformational states in the local framework (closed heads or circles), while the internal spatial relations among four states in either set of compared data were fixed. For illustration, we calculated the combined least-square (*LS*_*c*_) values between the two sets of open and closed arrow heads in Fig. 8A for all 24 combinations, and plotted the 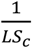 values in Fig. 8C. In the most probable combination based on the largest 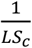 corresponds to E_x_, *C*_2_ to E_o_, *C*_3_ to I_o_, and *C*_4_ to I_x_, in which the values *θ*_i_ and *φ*_i_ and *Ω* are comparable between the corresponding conformational and structural states (Fig. 6D,E). These assignments are supported by the independent evidence that the probabilities of *C*_2_ and *C*_3_, which correspond to open states, vary with the concentration of Arg^*+*^ whereas those of *C*_1_ and *C*_4_, which correspond to occluded states, stay relatively constant (Fig. 9A).

The present study was performed under equilibrium conductions, which allowed us to determine straightforwardly the *K*ssvalues of four conformations from the dependence of their probabilities on the ligand concentration. The resulting values of *K*_D1_ through *K*_D4_ ranges from 25 to 69 μM (Table 1), statistically comparable with the previously reported overall *K*_D_ of 32 to 93 μM determined by ITC (Fang et al., 2007, Tsai et al., 2012). Thus, maneuvers such as introducing cysteine mutations and attaching the fluorophore to the AdiC molecule do not appear to have any marked impact on its affinity for Arg^*+*^. Such a finding is not particularly surprising because the chosen labeling part is on the surface of the protein such that it is not at, but external to, the ligand-binding site.

In summary, using a state-of-the-art fluorescence polarization microscopic system assembled in house, we have, for the first time, successfully tracked conformational changes in a single integral membrane protein molecule among four states that occur on angstrom and millisecond scales. The resolving power of this technique allowed us to relate the four conformations to known structure-function states, and to experimentally establish a fully determined quantitative model of 8 states required to account for the conformational energetics of a AdiC transport molecule with or without a ligand bound. This capability of resolving and tracking the conformational states of individual molecules forms the foundation for us to perform the kinetic study to acquire the necessary dynamic information for understanding the transporting mechanism and for creating an experiment-based 4D-model to quantitatively account for the complex spatiotemporal behaviors of a transporter molecule.

## Materials and Methods

### Materials

Detergent n-Dodecyl-β-D-Maltopyranoside (DDM) was purchased from Anatrace, 1-palmitoyl-2-oleoyl-glycero-3-phosphocholine (POPC) from Avanti Polar Lipid Inc., bifunctional rhodamine bis-((N-iodoacetyl)-piperazinyl)-sulfonerhodamine from Invitrogen (B10621), strep-tactin resin from IBA, cover slip (#1.5) and microscope slide glass from Fisher Scientific or VWR. Unless specified otherwise, all other reagents were purchased from Sigma, Thermo Fisher Scientific, or EMD Millipore.

### Cloning, Protein expression and purification

A double-stranded DNA fragment, synthesized by Integrated DNA Technologies (IDT), contains, from the N-terminus to the C-terminus, Avi tag for recognition by biotin ligase, a strep-tag, a linker (GGGSGGGS), the gene of AdiC of *E. coli*, a linker (GGGS), a thrombin protease recognition site, a C-termini strep-tag, and a stop codon, which was cloned into the pET28b vector. The removal of two native cysteines (C238A and C281A) and introduction of the double G188C and S195C cysteine mutations in helix 6a for attaching bifunctional rhodamine were carried out using the QuickChange technique and verified by DNA sequencing.

The AdiC protein was expressed in *E*.*coli* BL21(DE3) cells transformed with AdiC-gene-containing plasmids. The transformed cells were grown in Luria Broth at 37°C to an A_600_ of ∼1.0. Protein expression was induced with 0.5 mM iso-propyl β-D-thiogalactopyranoside (IPTG) at 22°C overnight. The cells were harvested and resuspended in a solution containing 100 mM NaCl, 50 mM tris-(hydroxymethyl)-aminomethane (Tris) titrated to pH 8.0, 1 mM phenylmethylsulfonyl fluoride (PMSF), 4 mM tris-(2-carboxyethyl)-phosphine (TCEP), 1µg/ml leupeptin and 1µg/ml pepstatin A. To extract membrane proteins, 40 mM DDM was added to the cell suspension; the conical tube containing this suspension was placed on a rotating device in a 4°C cold room for 2 - 3 hours. The cell suspension was then sonicated, the cell lysate was centrifuged at 12,000 x g for 20 min, and the resulting supernatant was loaded onto a gravity flow column packed with strep-tactin super-flow resin. The column was washed with a wash buffer (WB) containing 100 mM NaCl, 50 mM Tris titrated to pH 8.0, 2 mM TCEP, and 2 mM DDM. AdiC protein was eluted from the column using WB added with 10 mM desthiobiotin. The AdiC containing fractions were pooled and then concentrated with an Amicon Ultra concentrator (50K MWCO) before a further purification with a size-exclusion FPLC column (superdex 200 10/30, GE) equilibrated in WB.

### Labeling and reconstitution of AdiC into nanodiscs

Purified AdiC was mixed with POPC and the MSP2N2 membrane scaffold protein purified as described (Ritchie et al., 2009, Denisov et al., 2019) in a 1 : 1500 : 10 molar ratio of the AdiC dimer: POPC : MSP2N2. The mixture was incubated at 4°C for at least 2 hours. Nanodiscs were assembled during a dialysis of the mixture against a buffer containing 100 mM NaCl, 20 mM Tris titrated to pH 8.0, and 0.5 mM TCEP at 4°C overnight. After the dialysis, the nanodiscs were labeled with biotin using the BirA Enzyme (Avidity) following the protocol provided by the manufacture, and then mixed with bifunctional rhodamine in a 2 : 1 molar ratio of the AdiC dimer: bifunctional rhodamine, a ratio to maximize the chance of only one AdiC monomer in each dimer to be labeled; this mixture was incubated at room temperature for more than 4 hours. The remaining free dye was removed by size-exclusion column (a superose 6 10/30, GE) equilibrated with a solution containing 100 mM NaCl and 50 mM Tris titrated to pH 8.0. The final product of nanodiscs harboring labeled AdiC was aliquoted, flash frozen in liquid nitrogen and stored in a −80°C freezer.

### Sample preparation for data collection with polarization TIRF microscope

For adhesion of streptavidin, one side of a cover slip is exposed to 0.01% poly-L-Lysine solution for 1 hour, before being rinsed with distilled water and air dried. Prior to an experiment, a cover slip was attached, via thin transparent Scotch adhesive tapes placed on its left and right edges, to a microscopy slide, with the poly-L-Lysine-coated side of the cover slip facing the bottom side of the slide. Solutions were to be placed in the space between the two pieces of glass created by the adhesive tapes that acted as a spacer. The poly-L-lysine coated side of the cover slip was exposed to 5 mg/ml streptavidin (Promega) in this space for 15 min, and the remaining free streptavidin was washed away with a solution containing 50 mM HEPES titrated pH 7.5 and 100 mM NaCl.

An AdiC-containing nanodisc sample was diluted to 30 - 100 pM, estimated from an evaluation of the absorbance of the sample at 550 nm wavelength against the extinction coefficient of bifunctional rhodamine. The diluted sample was flowed into the space between the assembled cover slip and slide. After allowing a biotin-moiety covalently linked to the N-terminus and the streptavidin-binding tags linked to the N- and C-termini in each of AdiC subunit (i.e., totaling six available attachment points per AdiC dimer) to bind to streptavidin on the cover slip, the space was thoroughly washed to remove unattached AdiC with a solution (pH 5) containing 100 mM NaCl, 100 mM dithiothreitol (DTT, Fisher, BP172) and 50 mM acetic acid, without or with arginine at a specific concentration. DDT was used to scavenge oxygen to minimize its adverse impact on the fluorophore’s emitting intensity and lifetime.

### Fluorescence polarization microscope and intensity recording

As previously described, the fluorescence polarized microscope was built from a Nikon TIRF microscope (model Ti-E) (Lewis and Lu, 2019c). To produce an evanescent field at the surface of the sample coverslip, a 140 mW linearly polarized laser beam (532 nm) generated from a 500 mW laser (Crystalaser CL532-500-S) was directed to pass through a ¼ λ-plate, which transformed the linear polarization to circular polarization. After passing through a polarization-preserving, high numerical aperture 100X objective (Nikon Achromatic, NA = 1.49), the beam emerged at a 68° angle incident to the coverslip, the so-called critical angle that leads to TIR required for the formation of an evanescent field (Axelrod et al., 1984). The emission of polarized fluorescent light from individual fluorophores excited by the evanescent field was directed to a 50 : 50 non-polarizing beam splitter (Thorlabs CM1-BS013) after passing through the objective (Fig. 2), and then to a 540/593 nm bandpass filter (Semrock FF01-593/40-25) that prevents the propagation of excitation light. One resulting beam was further split by a glass (N-SF1) polarizing beam splitter (Thorlabs CM1 PBS251) along 0° and 90° and the other by a wire-grid polarizing beam splitter (Thorlabs WP25M-Vis) along 45° and 135°. These 4 emission intensity components, labeled as *I*_0_, *I*_45_, *I*_90_and *I*_135_, were individually directed onto 4 designated sectors in the CCD grid of an EMCCD camera (Andor iXon Ultra 897), where the four intensities from a given fluorophore appeared in the corresponding positions of the four sections.

Here, fluorescence intensities from individual bifunctional rhodamine molecules, each attached to helix 6A in AdiC, were collected with the microscope and captured every 10 ms with an EMCCD camera at the room temperature 22ºC. Following extraction of temporal information from the intensities with the changepoint analysis described below, we applied a Gaussian filter (with a corner frequency of 7.5 Hz) to all four intensity traces to reduce high-frequency noise, where the rise time was 22 ms. From these filtered intensities, we calculated angles as described below.

The experiments were performed in five separate occasions. Data collected among these separate collections are statistically comparable and were pooled together, resulting in sufficiently narrow distributions as illustrated in exhibited in Fig. 7. The width of the distributions reflects both technical and biological variations. Outlier data were excluded on the following basis. First, while fluorescence intensity is expected to vary among different polarization directions, the total intensity should be essentially invariant. Therefore, any recordings whose total intensity changes beyond what were expected for random noise are excluded. Second, for a given recording, at least 12 events are required to obtain a 95% confidence level for state identification, so any traces with less than 12 events are excluded. Third, for event detection and state identification, a signal to noise ratio greater than 5 is necessary for the required minimum angle resolution. Thus, any set of intensity traces with this ratio less than 5 are excluded. The sample sizes were estimated on the basis of previous studies (Lewis and Lu, 2019c, Lewis and Lu, 2019a, Lewis and Lu, 2019b) to yield sufficiently small standard errors of mean to obtain accurate estimate of the mean. Practically, the error bars are comparable to the sizes of the symbols of data as illustrated in Fig. 9A. The 95% confidence intervals are provided for all determined equilibrium constants in Table 1.

### System parameters

Theoretical aspects of the 3 channel polarized emission system have been previously described (Fourkas, 2001), which has been extended to and practically implemented with 4 channels (Ohmachi et al., 2012, Lippert et al., 2017, Lewis and Lu, 2019c). Briefly, the intensity of a given polarized component with angle Ψ, *I*_Ψ_, collected by the microscope’s objective from the emission of a fluorophore, which is excited by a circularly polarized TIRF field, is dependent on the fluorophore’s orientation. As such, *I*_ψ_is defined by the spherical coordinates *θ* and *φ* in a manner that

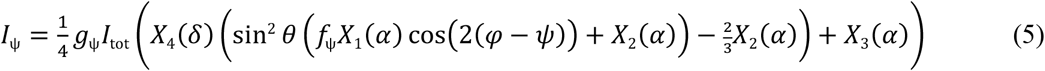

where the polarization angle *ψ*= 0°, 45°, 90° or 135°. The factor *f*_ψ_corrects for systemic reduction in the maximal achievable anisotropy of the light per channel *Ψ*. The coefficients *X*_1_, *X*_2_ and *X*_3_ correct for the incomplete collection of photons by a microscope objective with collection half-angle *α*:

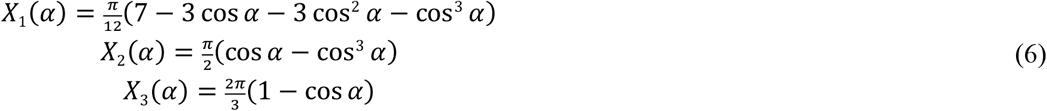

Ideally, when presented with a beam of non-polarized light, the system splits that beam into four of equal intensity. Small deviations from this theoretical equality are corrected by normalizing each intensity of a given channel (*Ψ*°) to that of the 90° channel chosen as the reference here:

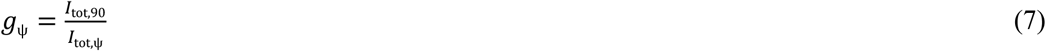

Besides those three types of system parameters, the coefficient *X*_4_ corrects for the fast diffusive motion (‘wobble’) of the probe relative to the attached protein, which is measured in terms of the half-angle *δ* of the wobble cone:

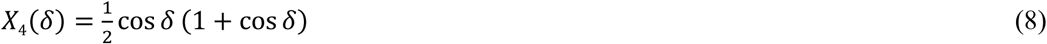

The parameter *δ* was experimentally estimated to be 22.5° in a separated macroscopic anisotropy study of the proteins as previously described (Lewis and Lu, 2019c).

Analytic solutions have been found for θ, φ and I_tot_ from Eq. 5 (Lewis and Lu, 2019c):

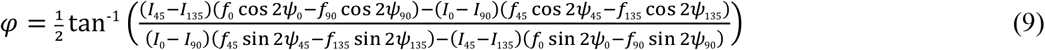

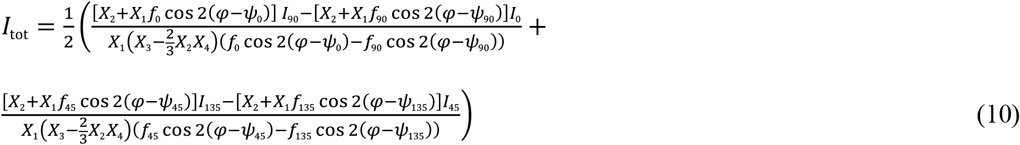

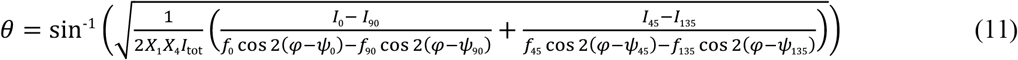

When *θ* is near 90°, the noise of intensities might prevent a solution of *θ*, in which case we simply set the *θ* values to 90°. We only analyzed data from the particles with no more than a few percent of such data points.

### Calibrations of the camera

Photons hitting the CCD chip of an EMCCD camera result in the release of individual electrons in accordance with the photo-electric effect. These electrons pass through multiple layers, at each of which they have a probability to cause the release of additional electrons, effectively amplifying the original signal with a gain (*G*) (Lidke et al., 2005, Heintzmann et al., 2016). The relationship of the *N* photons released over a given interval and the recorded intensity *I* is expressed as

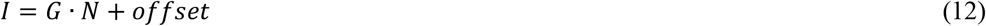

As an inherent feature of EMCCD cameras, the offset is the intensity recorded by the camera when the shutter is fully closed. These parameters are estimated by analyzing the relationship between photon intensities recorded at multiple laser intensities versus the corresponding SNR. For photon count *N*, the standard deviation *σ* due to shot noise is given by the square root of *N*. In addition, EMCCD cameras add an additional multiplicative noise that effectively scales the *σ* due to shot noise by a factor of √2 relating the SNR to *N* by the expression

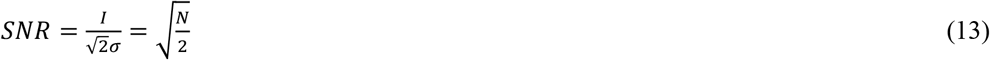

Substituting Eq. 13 into Eq. 12 yields

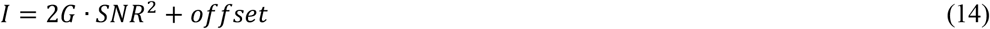

On a plot of *I* versus SNR^2^, the slope is twice the gain whereas the y-intercept is the offset. For our system, the value of the gain was found to be 146 and that of the offset was 220.

Furthermore, when the signal from an interested fluorophore is split from the background signal, the combined intensity signal recorded is related to *N* by the following relation

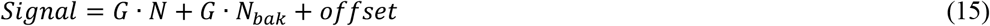

rearranged to

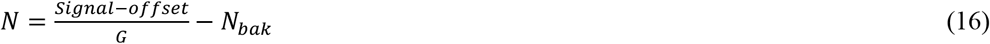

where *N*_𝑏𝑎k_ is the number of photons underlying the background signal, which needs to be subtracted. The quantum efficiency (QE) of the Andor Ixon emccd camera is estimated as ∼0.95 in the visible light range, meaning that 95% of photons contacting the ccd chip are detected. Therefore, the actual photon count is calculated as N_det_ = N / QE, and the effective SNR is also scaled accordingly.

### Detection of event transitions

The photon release rate changes when the orientation of the fluorophore is changed. To detect these transitions within the measured polarized intensities, we adopted a version of the changepoint algorithm (Chen and Gupta, 2001, Beausang et al., 2008). Such a process was based on calculating a log likelihood ratio over a period of time to determine the maximal ratio that identified the point where the change of photon-release rate occurred, i.e. the time at which the fluorophore transitions from one orientation to another. This method has previously been applied to analyzing the photon-arrival time captured on a continuous basis with photon-multiplier-based multi-channel recordings (Beausang et al., 2008), which was adapted for analyzing photons collected over a fixed time interval with an EMCCD camera(Lewis and Lu, 2019c).

When a camera is used as a detector, photons emerging from each channel are effectively binned over each frame. A series of consecutive *k* frames with a constant exposure time (Δ*t*) is expressed as

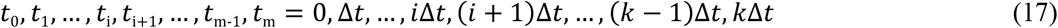

For a given frame *i*, the intensity *I*_i_ is defined by the rate of photon release:

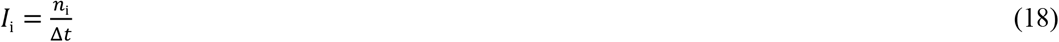

where *n*_i_ is the number of photons release within frame *i*. The cumulative distribution, *m*_j_, is built by adding the number of photons for the successive time frames:

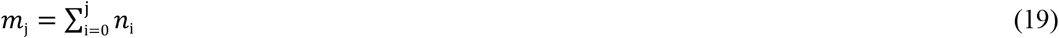

If during an interval *T* the rate changes from *I*_1_ to *I*_2_ at the time point *τ* = *f*Δ*t*, and the number of emitted photons prior to this change is *m* (Eq. 19) and *N - m* after the change, then the likelihood ratio of a transition occurring at frame *f* in the log form is described by

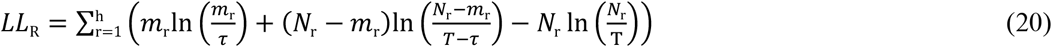

where *h* equals 4, the number of emission channels in the system. We set the threshold of significance for *LL*_*R*_ at the level that limits the false positive events to 5% on the basis of simulation studies for the corresponding levels of SNR, and the resulting false negative events were about 1%.

The program is started by identifying one transition over the entire trace. If a change-point X was identified, it would then search for additional transitions between the start of the trace and point X and between X and the end. This iterative search with successively shortened stretches continued until no more transitions were identified.

### State identification

Following the detection of intensity transition timepoints using the changepoint method and subsequent calculation of angles, the states of individual events, each demarcated by two consecutive transition timepoints, were identified on the basis of the angles *θ* and *φ* together (Lewis and Lu, 2019c). The resulting four conformational state distributions are denoted as *C*_1_ – *C*_4_. This identification of the states is done by using a ‘nearest-neighbor’ method in a cartesian coordinate system. As such, the x,y,z, values were calculated from the corresponding *θ* and *φ* values, along with a unity *r* in accordance with the transformation relations between the spherical and cartesian coordinate systems:

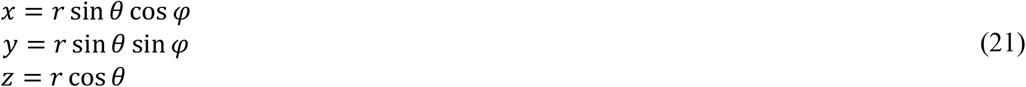

Given the unity *r*, which carries no information regarding spatial orientation, for all cases, x,y,z in all cases would always be on a unit sphere and fully encode the orientation information specified by *θ* and *φ*.

States must be indexed so that they are always numerated in the same sequential order among individual molecules, a prerequisite for correctly relating the states identified here to those identified crystallographically. This indexing is based on the shortest, if not straightest, path distance between the first and last states, *C*_1_ and *C*_M_ (where *M* = 4 in our case) as they detour through the remaining states (Fig. S1B). This path length is calculated as the sum of the accumulative distances between two adjacent states, denoted as *d*_tot_:

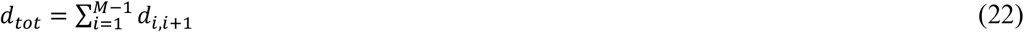

where the number 1 indicates a chosen starting state and the distance *d*_i,j_ between states indicated by positions *i* and *j* in a cartesian coordinate system is defined as:

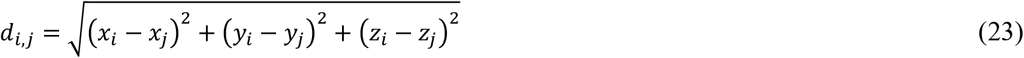

If the states were located along a perfect line, the total path distance would equal the distance between states 1 and M, i.e. *d*_1,M_, where *d*_1,M_ = *d*_tot_. However, if they were not on a line, *d*_tot_ would be greater than *d*_1,M_ by *Δd* such that

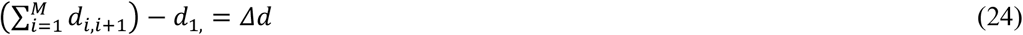

Upon calculating *Δd* for each of the 24 possible sequences relating the four states, the shortest path can be found on the basis of the smallest value of *Δd*, which has two solutions with equal path length, i.e., 1-2-3-4 versus 4-3-2-1 (Fig. S1C versus S1D). The sequence of *C*_1_-*C*_2_-*C*_3_-*C*_4_ shown Fig. S1C was consistently chosen.

### Transformation of the laboratory framework to a local framework

As explained in the text, individual molecules do not have the same orientation. Thus, a direct comparison among them requires all molecules be rotated from the laboratory frame of reference into a common local frame of reference (Fig. S1A). Here, the x,y-plane is defined by the vectors representing *C*_1_ (**V**_**1**_) and *C*_4_ (**V**_**4**_) and the x-axis is defined by **V**_**1**_. The z-axis is perpendicular to the x,y-plane. The **x, y** and **z**-axes, presented as unit vectors, are then defined by:

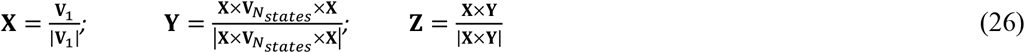

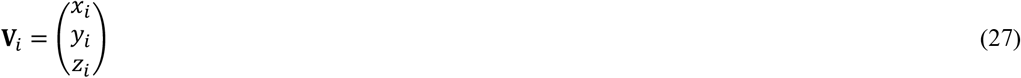

*θ* and *φ* in the local frame of reference are then calculated as:

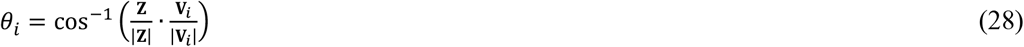

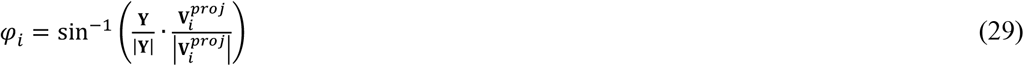

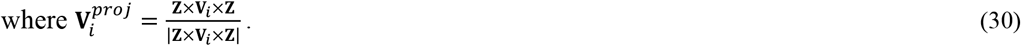

### Calculation of the direct angle change Ω between two states

The direct angle change *Ω*_i,j_ between two states *i* and *j*, as represented by the vectors **V**_i_ and **V**_j_ defined above, can be calculated from the relation.

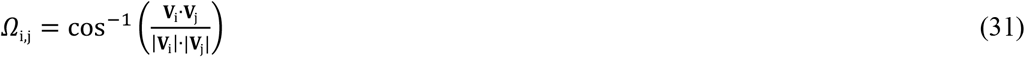

## Acknowledgement

This study was supported by the grant DK125521 from the National Institute of Diabetes and Digestive and Kidney Diseases.

## Data availability

A zipped source-data file for all figures is provided.

## Supplementary Figure Legends

**Figure S1.**
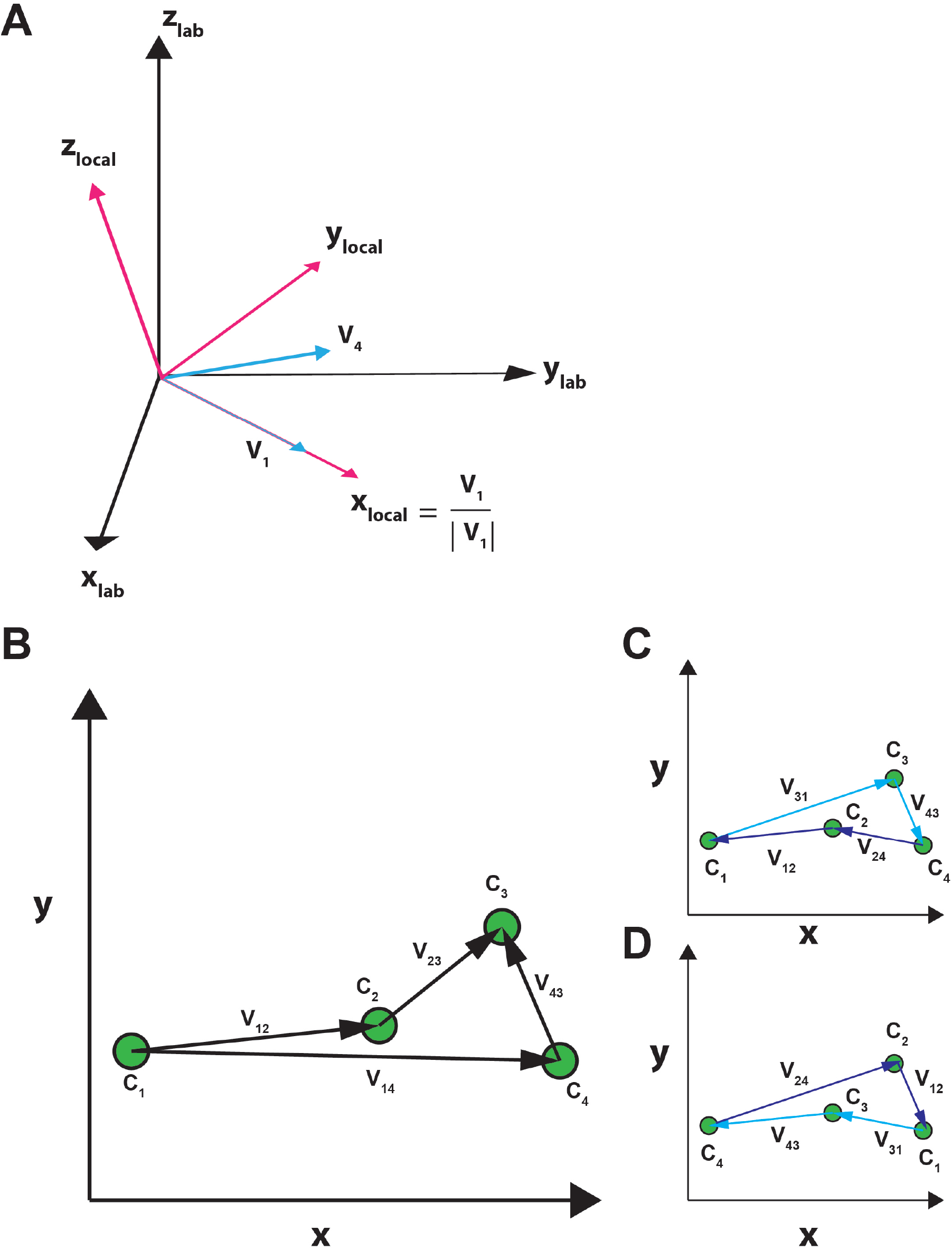
Transformation between the laboratory and local coordinates and assignment of states. (A). Coordinate transformation. In the laboratory frame of reference (colored black), the z_lab_-axis is defined as parallel to the microscope objective and the x_lab_,y_lab_-axes are defined within the plane of the sample glass, perpendicular to the objective and the z_lab_-axis. Coordinates are transformed to the local frame of reference defined by the vectors **V**_1_ and **V**_4_ that represent the orientations of *C*_1_ and *C*_4_, respectively, such that the new x-axis becomes parallel to the vector **V**_1_ (Eq. 26, Methods), the new y-axis becomes perpendicular to the new x-axis in the plane defined by **V**_1_ and **V**_4_, and the new z-axis becomes perpendicular to the new x,y-plane. (B-D) State assignment. Among four states, there are 24 unique serial relations. The path between a given pair of states *C*_i_ and *C*_j_ are indicated by a vector **V**_i,j_ between them (B). To ensure the same sequence among the four states is assigned in all particles analyzed individually, states *C*_1_ through *C*_4_ are aligned such that the calculated distance among them, defined by |**V**_12_|+|**V**_23_|+|**V**_34_| is shortest. Thus, their combined distance is closest to the length of the direct path between *C*_1_ and *C*_4_ defined by the length of **V**_14_ (Eq. 24, Methods). This operation should, in principle, yield two solutions with inverted sequences: *C*_1_-*C*_2_-*C*_3_-*C*_4_ (**c**) versus *C*_4_-*C*_3_-*C*_2_-*C*_1_ (C). The sequence of *C*_1_-*C*_2_-*C*_3_-*C*_4_ in C was consistently chosen.

## Description of Supplementary Video 1

**Video 1. A** 7.2 second long video of the four channel emission intensities captured on the EMCCD, ordered top to bottom as *I*_0_, *I*_45_, *I*_90_ and *I*_135_. These intensities are also shown in Fig. 3A. Running traces of the intensities integrated from the original movies are displayed in the middle section, and the angle traces, *θ, φ* and *Ω* (with *C*_1_ as a reference) calculated from the intensities are shown in the right section.

